# Envirotyping Facilitates Understanding of Genotype × Environment Interactions and Highlights the Potential of Stay-green Traits in Wheat

**DOI:** 10.1101/2025.03.11.642575

**Authors:** Asad Amin, Jack Christopher, Clayton Forknall, Mark Cooper, Bethany Macdonald, Brian Collins, Kai Voss-Fels, Lee Hickey, Karine Chenu

## Abstract

Better understanding genotype by environment interaction (G×E) can help breeding for better adapted varieties. Envirotyping for environmental water status was applied to assist interpretation of G×E interactions for wheat yield in multi-environment trials conducted in drought-prone Australian environments. Genotypes from a multi-reference parent nested association mapping (MR-NAM) population were tested in 10 trials across the Australian wheatbelt. Genotype yield and phenology were measured in all trials, while traits associated with the stay-green phenotype were assessed for a subset of 5 trials. Envirotyping was conducted by characterizing water stress experienced by genotypes at each trial using crop modelling. Envirotyping facilitated the understanding of G×E interactions by explaining 75, 67, and 66% of the genotypic variance for yield in severe water-limited (ET3), mild terminal water-stress (ET2), and water-sufficient (ET1) environments, respectively.

Yield and stay-green were negatively correlated with flowering time in most trials. However, when focusing on genotypes flowering at similar times within a trial, no significant correlation was found between yield and flowering. Importantly stay-green traits remained significantly correlated with yield. Stay-green traits such as delayed onset of senescence and slower senescence rate benefited yield by 0.2 to 1.1 t ha^-1^ across environments, highlighting the breeding potential for stay-green traits in both water-sufficient and water-limited environments. Hence, sustaining green leaf area during grain filling helped to enhance yield. Envirotyping to better understand G×E interactions for yield, coupled with screening for traits exhibiting superior adaptive mechanisms, are powerful assets in assisting plant breeders to select more effectively drought adapted genotypes.

**Highlights:**

Genotype × environment interaction for yield could be reduced with envirotyping.
Envirotyping enabled a 75% gain in genotypic variance in severe drought conditions.
Envirotyping clarified breeding potential of stay-green traits across environments.
Stay-green has potential to enhance wheat yield across diverse environments.

## 1 Introduction

The development of agricultural systems and plant breeding is continually evolving (Cooper et al., 2014). Crop improvement in breeding programs is limited by complex Genotype by Environment (G×E) interactions associated with genotype plasticity (trait response to environment; Basford and Cooper 1998). This genotype plasticity varies with the timing, severity, and duration of stress factors during the physiological development of plants (Hammer et al., 2006; Fischer, 2011; Vadez et al., 2024; Gawinowski et al., 2025).

Considerable variation in the climatic and soil conditions exist across agricultural regions, such as across the Australian wheatbelt (Chenu et al., 2013). In addition, these conditions are subject to strong inter-annual changes (Potgieter et al., 2002; Ray et al., 2015), which are evolving with climate change (e.g. IPCC, 2019; Ababaei and Chenu, 2020). Such environmental variation leads to G×E interactions in farmers’ fields and breeders’ trials. However, broad-scale field testing of all G×E combinations in the target population of environments (TPE) is beyond the capacity and resources of most breeding programmes, even when considering only a relatively limited number of genotypes and environments in multi-environment trials (METs) (Cooper et al., 2014). In this context, characterizing the environment experienced by crops with ‘envirotyping’ can assist breeding programs (Cooper et al., 2021; Resende et al., 2022) as this helps to (i) reduce G×E interactions within environment types (Löffler et al., 2005; Chenu et al., 2011; Chenu, 2015), (ii) better understand the TPE (Chapman et al., 2000b; Chenu et al., 2013; Watson et al., 2017) and (iii) weight trial data to reduce undesirable influences of bias when samples of environments within a MET misrepresent the environmental composition of the TPE (Podlich et al., 1999, Messina et al., 2023).

Environmental characterization for wheat crops across the Australian wheatbelt has identified four representative drought environment types (ETs), defined by specific water-stress patterns over the crop cycle (Chenu et al., 2013). This approach helped to explain occurrences of G×E interaction for different genotypes (Chenu et al., 2011). In this TPE, where drought and heat stress are common during the grain filling period (Chenu et al., 2013; Ababaei and Chenu, 2020), early water saving associated with transpiration efficiency or traits such as deep rooting and stay-green that allow the crop to function for longer, have shown promise (e.g. Manschadi et al., 2006; Casadebaig et al., 2016; Christopher et al., 2016; Collins et al., 2021; Borrell et al., 2023; Shazadi et al., 2024). Stay-green in particular is an interesting phenotype that can be easily measured for large numbers of genotypes (e.g. Lopes and Reynolds, 2012; Christopher et al., 2014; Chapman et al., 2021). Genotypes with stay-green are known to retain green leaves for longer after flowering than standard genotypes allowing plants to carry out photosynthesis for longer during the grain-filling period (Thomas and Ougham, 2014). While the stay-green phenotype observed under terminal drought may result from water saving during early development (e.g. small canopy resulting in low water loss; Borrell et al., 2014) or greater access to soil water (e.g. deeper root system; Manschadi et al., 2006; Christopher et al., 2023), traits characterizing the stay-green phenotype have been found to correlated with yield in both water-sufficient and water-limited environments (Christopher et al., 2016). However, in some drought-prone environments, the stay-green phenotype has also been associated with yield loss (Chairi et al., 2020), as a longer crop duration can lead to greater risk of terminal drought exposure (Chenu et al., 2013; Vadez et al., 2024).

This paper aims to (i) assess how environment characterization can help to untangle G×E interactions for yield in a large MET dataset with diverse genetic variation and trials from across the Australian wheatbelt, (ii) investigate the relationship between stay-green traits and yield in different ETs, and (iii) explore the plasticity of stay-green traits across environments. To this end, 10 trials were conducted across the Australian wheatbelt with 1,819 wheat lines from a multi-reference parent nested association mapping (MR-NAM) population including parental lines with enhance stay-green (Richard, 2018). Envirotyping and GxE analysis for yield was conducted across all trials, while stay-green plasticity and its correlation with yield were investigated in a subset of five trials where normalized difference vegetation index (NDVI) was measured over time during grain filling.

## 2 Material and methods

### 2.1 Overview

Yield and flowering time were measured on subsets of the MR-NAM population tested at 10 field trials located across the Australian wheatbelt (Table 1, Fig. 1). The MR-NAM population was developed with a constrained range of plant height and phenology within each family (Richard, 2018). Genotypic correlations between trials were estimated via a mixed model MET analysis. Crop water-stress patterns were simulated for four reference lines at each trial with the APSIM-Wheat crop model (Holzworth et al., 2014; Zheng et al., 2015) to characterize their drought environment types (ETs). These trial ETs were included in the factor analytic (FA) based linear mixed model to assess how envirotyping may improve our understanding of G×E in the studied MET dataset.

**Fig. 1.**
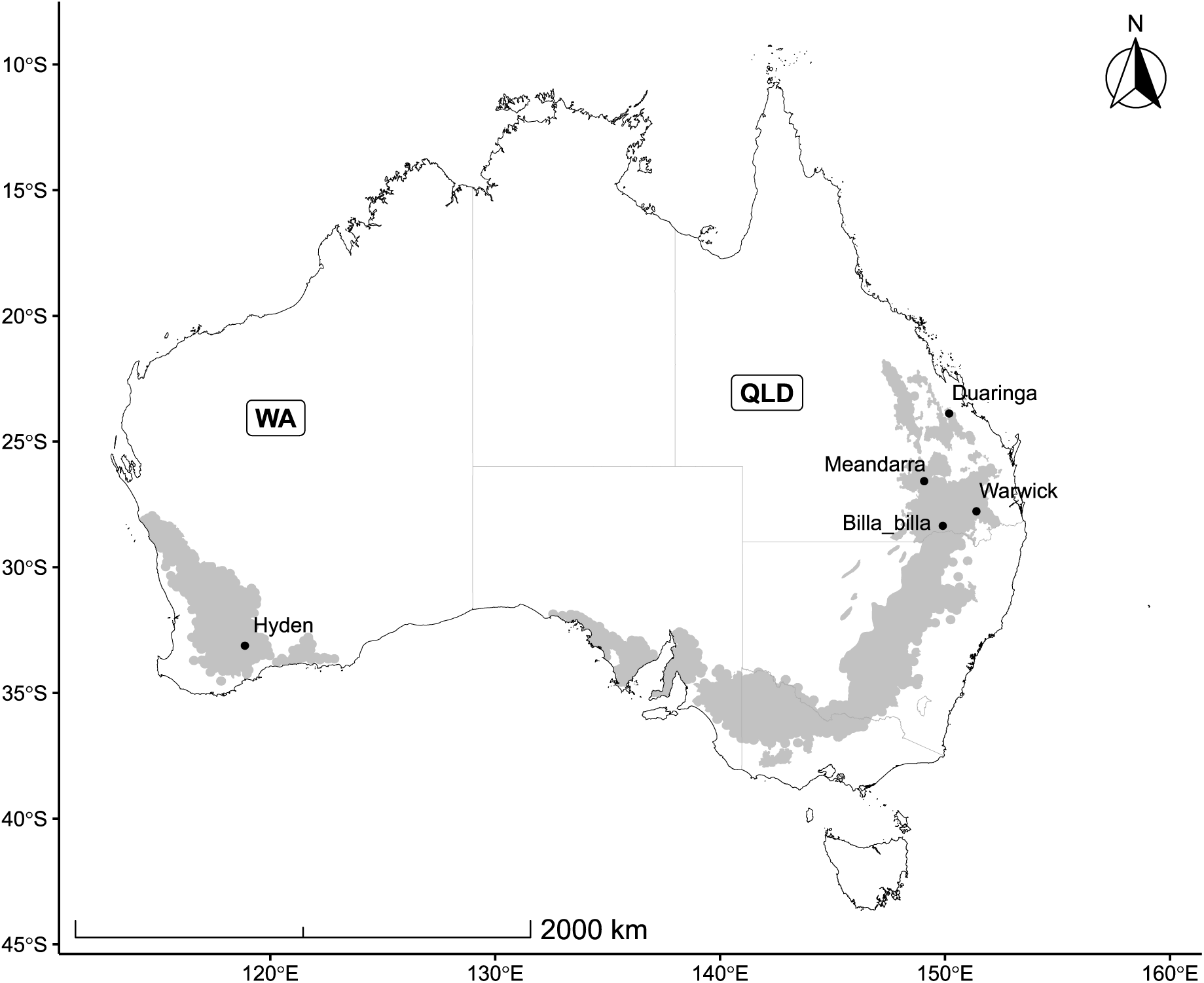
Map of the five locations where the 10 trials were conducted across the Australian wheatbelt (grey area). Labels indicate the location name (i.e. Warwick, Billa Billa…) and bold acronyms indicate state names (i.e. Western Australia (WA), Queensland (QLD)).

**Table 1.** Trial characteristics with the trial name, trial location, state, type of data collected (yield with or without monitoring of stay-green (SG)), sowing date, nitrogen fertilization applied at sowing (as urea for Queensland (QLD) and as nitrate in West Australia (WA)), number of days from sowing to flowering (DTF) and to maturity (DTM) for the reference parent Mace, number of genotypes tested in each trial (Lines), observed range of days to flowering (ORDF) for all studied genotypes, number of genotypes flowering around the population median i.e. within ‘selected range of days to flowering’ (SRDF), SRDF, drought environment type (ET), the trial-mean value of yield (Yield), maximum greenness around flowering (Nmax), start of senescence (OnS in °Cd since flowering), mid of senescence (MidS in °Cd since flowering), end of senescence (EndS in °Cd since flowering), indicator of the senescence rate (SR), and the stay-green integral (SGint) calculated within each trial. Each trial was named as location (i.e. WAR (Warwick), DUA (Duaringa), MEA (Meandarra), BIL (Billa-Billa), HYD (Hyden)), year (2014-2018), and irrigation status (rf (rainfed), and ir (irrigated)).

| Trial | Location | State | Data type | Sowing date |  | DTF<br>(d) | DTM<br>(d) | Lines | ORDF<br>(day) | Lines<br>within<br>SRDF | SRDF<br>(day) | ET | Yield<br>(t ha<sup>-1</sup>) | Nmax | OnS<br>(°Cd) | MidS<br>(°Cd) | EndS<br>(°Cd) | SR | SGint |
| --- | --- | --- | --- | --- | --- | --- | --- | --- | --- | --- | --- | --- | --- | --- | --- | --- | --- | --- | --- |
| WAR14rf | Warwick | QLD | Yield/SG | 10/07/2014 | 120 | 96 | 123 | 802 | 81-104 | 714 | 88-93 | 1 | 7.25 | 0.76 | 191 | 318 | 536 | 4.34 | 217 |
| WAR15rf | Warwick | QLD | Yield/SG | 11/06/2015 | 120 | 111 | 143 | 975 | 103-130 | 668 | 112-115 | 1 | 6.58 | 0.80 | 230 | 441 | 744 | 6.77 | 515 |
| WAR15ir | Warwick | QLD | Yield | 11/06/2015 | 120 | 111 | 143 | 162 | NA | NA | NA | 1 | 7.16 | NA | NA | NA | NA | NA | NA |
| DUA15rf | Duaranga | QLD | Yield | 15/05/2015 | 120 | 78 | 115 | 191 | NA | NA | NA | 1 | 3.23 | NA | NA | NA | NA | NA | NA |
| MEA15rf | Meandarra | QLD | Yield | 19/05/2015 | 120 | 99 | 140 | 191 | NA | NA | NA | 1 | 3.79 | NA | NA | NA | NA | NA | NA |
| WAR16rf | Warwick | QLD | Yield/SG | 22/07/2016 | 120 | 96 | 128 | 866 | 78-104 | 559 | 81-83 | 2 | 4.87 | 0.75 | 228 | 379 | 577 | 9.03 | 438 |
| WAR17rf | Warwick | QLD | Yield/SG | 27/05/2017 | 120 | 100 | 131 | 679 | 93-118 | 630 | 98-107 | 3 | 3.83 | 0.84 | -26.9 | 127 | 387 | 5.38 | 323 |
| WAR17ir | Warwick | QLD | Yield/SG | 27/06/2017 | 120 | 100 | 131 | 210 | 100-115 | 178 | 103-108 | 1 | 6.28 | 0.84 | 31.1 | 223 | 586 | 6.21 | 411 |
| BIL18rf | Billa-Billa | QLD | Yield | 24/05/2018 | 120 | 101 | 136 | 665 | NA | NA | NA | 3 | 2.34 | NA | NA | NA | NA | NA | NA |
| HYD18rf | Hyden | WA | Yield | 02/05/2018 | 95 | 114 | 151 | 161 | NA | NA | NA | 2 | 2.03 | NA | NA | NA | NA | NA | NA |

In five of the 10 studied trials, stay-green phenotypes were estimated by using normalized difference vegetation index (NDVI) measured over time during grain filling (Fig. 2; Christopher et al., 2014; 2016). Weekly NDVI measurements from 300°Cd before flowering onwards, were fitted to a logistic curve, and features of the curve enabled the estimation of stay-green traits. The value of these traits for explaining yield variation in the MET dataset was estimated based on correlations with yield.

**Fig. 2.**
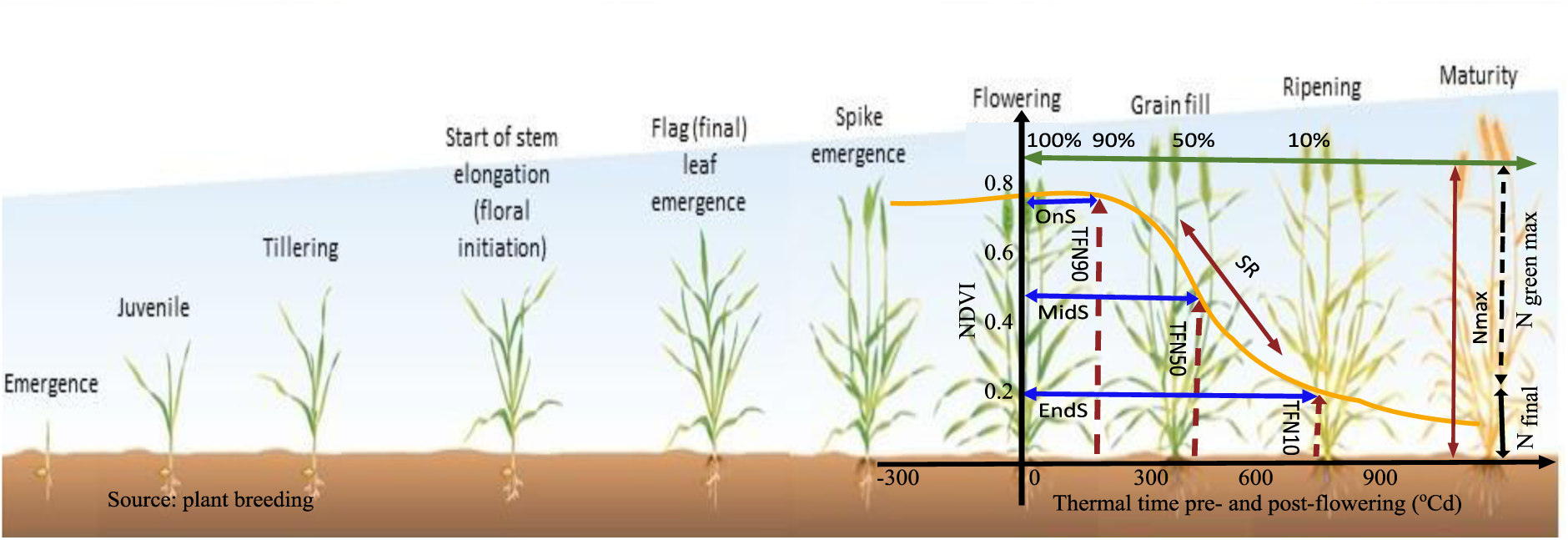
Stay-green traits based on the change in normalized difference for vegetative index (NDVI) over time: start of senescence (OnS), mid of senescence (MidS), end of senescence (EndS), indicator of the senescence rate (SR), maximum greenness around flowering (Nmax) and the timing TFN90, TFN50, and TFN10 (in degree days), which correspond to OnS, MidS, and EndS, respectively. Figure adapted from Christopher et al. (2014).

Genotypic correlations of flowering time with (i) stay-green traits, and (ii) yield were also quantified. For each trial, the yield advantage of stay-green traits was estimated by the yield difference between genotypes having the top and bottom 5 percentile values for each stay-green trait.

### 2.2 Genotypes

Three Australian spring-habit wheat (*Triticum aestivum* L.) cultivars (Suntop, Mace, and Scout) were used as reference parents to develop a MR-NAM population (Christopher et al., 2021; Richard, 2018). Suntop, Mace, and Scout are modern cultivars widely grown in the Australian eastern, western, and southern wheat production regions, respectively. These reference parents are known to have resistance to leaf and stem rust and other diseases (www.dpi.nsw.gov.au). Eleven founder lines with some adaptative traits of interest were used (Dharwar-dry, Drysdale, EGA-Gregory, Face10-16, Ril114, SB062, Seri-M82, Westonia, Wylie, ZWB10-37, and ZWW10-50). Dharwar-dry, Drysdale, and SeriM82 are known to have deep and dense root systems (Manschadi et al., 2006; Christopher et al., 2008) with improved stay-green phenotype (Olivares-Villegas et al., 2007; Christopher et al., 2008; Manschadi et al., 2010). Drysdale is an Australian variety known for improved transpiration efficiency (Condon et al., 2004; Rebetzke et al., 2009; Tausz-Posch et al., 2012; Fletcher et al., 2018; Collins et al., 2021). SB062 is a new breeding line developed by International Maize and Wheat Improvement Center (CIMMYT) with improved tolerance for extreme weather conditions, which also comes with low canopy temperature and more water-soluble carbohydrates (Dreccer et al., 2009; Ullah and Chenu, 2019; Chenu and Oudin, 2019; Ullah et al., 2023). CIMMYT lines (ZWB10-37, and ZWW10-50) were selected from a joint adventure of Australia and India in CIMMYT-Australia and ICARDA Germplasm Evaluation (CAIGE; Trethowan et al., 2024) based on their high yield performance in Australian environments. EGA Gregory and Wylie are known to have multiple disease resistance, like root lesion nematodes (*Pratylenchus thornei*) and Fusarium crown rot resistance, respectively. These cultivars were selected from the Queensland Department of Agriculture and Fisheries (QDAF) breeding program (Queensland Wheat Variety Guide, 2014; Zheng et al., 2014). Westonia is an Australian cultivar known for tolerance to acidic soils, manganese, and aluminium toxicities (Khabaz-Saberi et al., 2010). Finally, RIL114 was selected for its high level of grain dormancy and pre-harvest sprouting tolerance (Hickey et al., 2009).

The MR-NAM development facilitated genome reshuffling throughout the crossing and development of recombinant inbred lines (RILs) with four generations of self-fertilisation. The MR-NAM had 6 Mace derived families (Ma-NAM), 6 Scout derived families (Sc-NAM), and 10 Suntop-derived families (Su-NAM) as previously described (Christopher et al., 2021; Richard, 2018).

A total of 1,819 lines of the MR-NAM were tested in the Australian wheatbelt, with the aim to develop a new gene pool for improving drought tolerance in modern elite Australian cultivars. Different subsets of the MR-NAM were tested in different trials based on their level of adaptation in each region.

### 2.3 Field trials

Ten trials were conducted at five locations across the Australian wheatbelt (Tables 1, 2; Fig. 1). Most of the experiments were rain-fed (rf). Irrigation (ir) was applied in some trials to have some water-sufficient environments. Two trials were irrigated (17 to 26 mm) pre-flowering to reduce the soil moisture deficit at this crucial plant growth stage. Soil type varied from very deep clays to sandy soils.

**Table 2.**
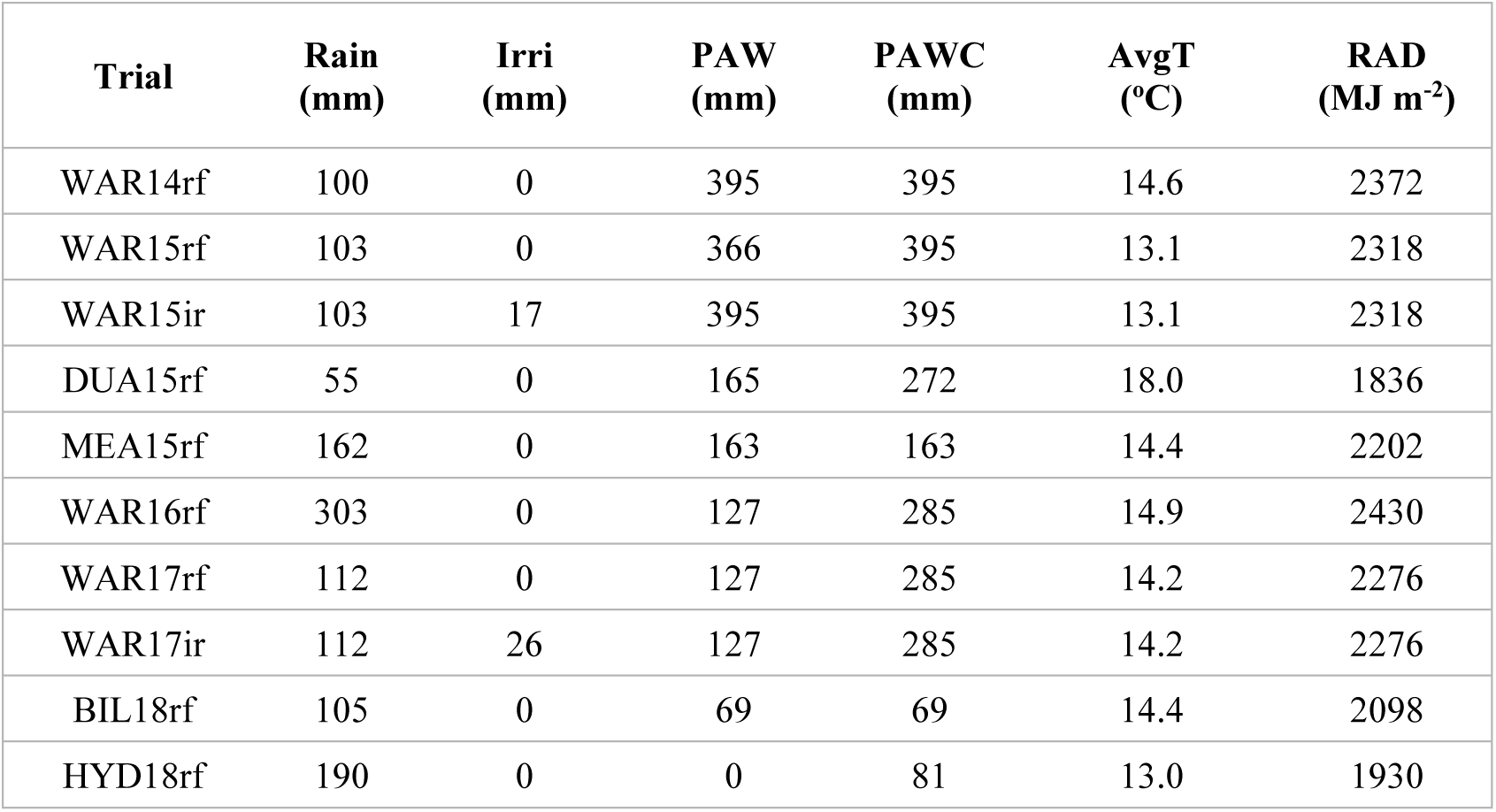
Trial environmental characteristics with the trial name, cumulative in-season rainfall (Rain, mm), pre-flowering irrigation (Irri), plant available water (PAW) of the soil at sowing, plant available water capacity (PAWC), average daily temperature from sowing to maturity (AvgT), cumulative radiation from sowing to maturity (RAD). Each trial was named as location (i.e. WAR (Warwick), DUA (Duaringa), MEA (Meandarra), BIL (Billa-Billa), HYD (Hyden)), year (2014-2018), and treatment (rf (rainfed), and ir (irrigated)).

Plants were cultivated for yield in 2 × 4 m plots with a row spacing of 25 cm and a target population density of 100 plants m^−2^. The area harvested was 2 × 4 m (8 m^2^), which included the outside rows (all 5 rows of each plot). Plot size at one of the first trials (WAR14rf) was smaller, 2 × 3 m, to accommodate the large number of genotypes. Pre-sowing soil moisture was measured, and soil tests were performed prior to planting to estimate soil nitrogen (N) and phosphorus (P) concentrations. Weeds and diseases were controlled as necessary. No crop failure was reported, but poor establishment occurred in two trials (WAR15ir and BIL18rf) for 2015 and 2018.

### 2.4 Stay-green traits

For 5 trials conducted at Warwick (QLD), a hand-held Green-seeker model 505 (NTech Industries, Ukiah, CA, USA) was used to measure the normalized difference vegetative index (NDVI) weekly from 300°Cd before flowering to 1500°Cd after flowering (Table 1). NDVI measurements were based on the variations in near-infrared and red reflectance for each plot. NDVI values from 300°Cd before flowering onwards were fitted by a logistic function (Equation 1) to estimate the stay-green traits for each genotype in each plot similarly as described by Christopher et al. (2014, 2021):

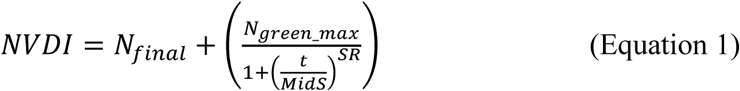

where *N*_*final*_ is NDVI value for fully senesced crops, *N*_*green*_*max*_ is the difference in maximum and final NDVI values (*ie*. *N*_*green*_*max*_ = *N*_*max*_ − *N*_*final*_ ), *t* is thermal time starting before flowering of the plot and expressed in °Cd centred at flowering, *MidS* is the thermal time from flowering to 50% of N_green_max_, and SR is an indicator of the senescence rate.

To enable proper fits to the data, an extra NDVI value corresponding to the final NDVI of the fully senesced crop was added after maturity, after the last measurement date. The NDVI value of this extra point was estimated by the minimum between 0.15 and the minimum NDVI recorded for the plot considered. Six stay-green traits were estimated based on Equation 1 (Fig. 2). These included the onset (OnS), mid (MidS), and end (EndS) of senescence, which are the thermal time from flowering with 90%, 50%, and 10% of N_green_max_, respectively. The initial value of NDVI indicated as maximum value of greenness (Nmax) and indicator of senescence rate (SR) in the plants was estimated from Equation 1. The stay-green integral (SGint) was calculated as the integral from 300°Cd before flowering to after flowering 1500°Cd, and corresponds to the area under the curve for this period between pre-flowering and maturity. SGint hence accounts for the overall crop greenness during the period considered. In addition to stay-green traits, yield was measured at harvest, and time to flowering was captured per plot in each trial.

### 2.5 Crop simulations and characterization of the water-sufficient and water-limited environment types

The Agricultural Production Systems Simulator (APSIM) wheat model (Holzworth et al., 2014; Zheng et al., 2015) was used to simulate the water deficit patterns at the studied field trials (Chenu et al., 2011; 2013) for four commercial varieties with contrasting maturity (Suntop, Mace, Hartog, and Gregory). Daily weather data were collected for nearby meteorological stations from the SILO based dataset (http://apsrunet.apsim.info; Jeffrey et al., 2001) for all but one trial (BIL18rf), where data were recorded on-site. Soil profiles developed by the Agricultural Production Systems Research Unit (APSRU) were sourced from the APSoil database (http://www.apsim.info/Products/APSoil.aspx) and selected based on similarities with local soil types. For trials at Warwick, soil information was available for the actual trials. As in previous studies (e.g. Chenu et al., 2011; Christopher et al., 2016), to best simulate the growth and development of the tested cultivars at each trial, an APSIM parameter (thermal time to floral initiation, ‘tt_floral_initiation’) was tuned so that simulated flowering time matched the mean observed flowering time (Fig. S1). In addition, soil moisture at sowing was optimized to decrease the deviation between observed and simulated yield. One unique soil moisture at sowing was used per site and year (i.e. the APSIM parameters for soil characteristics were the same for all genotypes and treatments (irrigation or rainfed) at each site).

Yield outputs of the crop model approximated adequately observed yield (Fig. 3), and the calibrated model was used to generate daily patterns of water-deficit index (Fig. 4). The water deficit index (‘supply-demand ratio’) represents the degree to which the soil water extractable by the roots (‘water supply’) is able to match the potential transpiration of the canopy (‘water demand’; Chenu et al., 2013). For each simulated cultivar and trial, the deficit index was centred at flowering and averaged every 100°Cd between emergence to 450°Cd after flowering, after which senescence greatly reduced plant transpiration and can thus lead to an ‘artificial’ increase of the water deficit and stress. Each water-deficit pattern was assigned to one of the four main drought environment types (ET1 - ET4) occurring in the Australian wheatbelt (Chenu et al., 2013) based on the minimum sum of squared differences for the trial water-deficit pattern compared with the representative water-deficit pattern of ET1 - ET4 (Fig. 4). Briefly, ET1 corresponds to no or minimum water deficit. ET2 represents mild deficit starting after flowering and relieved during the grain filling period. ET3 was characterized by more severe water deficit starting before flowering and relieved during grain filling (Chenu et al., 2013). None of the 10 studied trials were classified as ET4, which corresponds to severe water deficit not relieved during grain filling. In the studied trials, ETs did not vary much across the different maturity types tested, so that the ET for Mace was considered as the representative ET for each trial in the analysis.

**Fig. 3.**
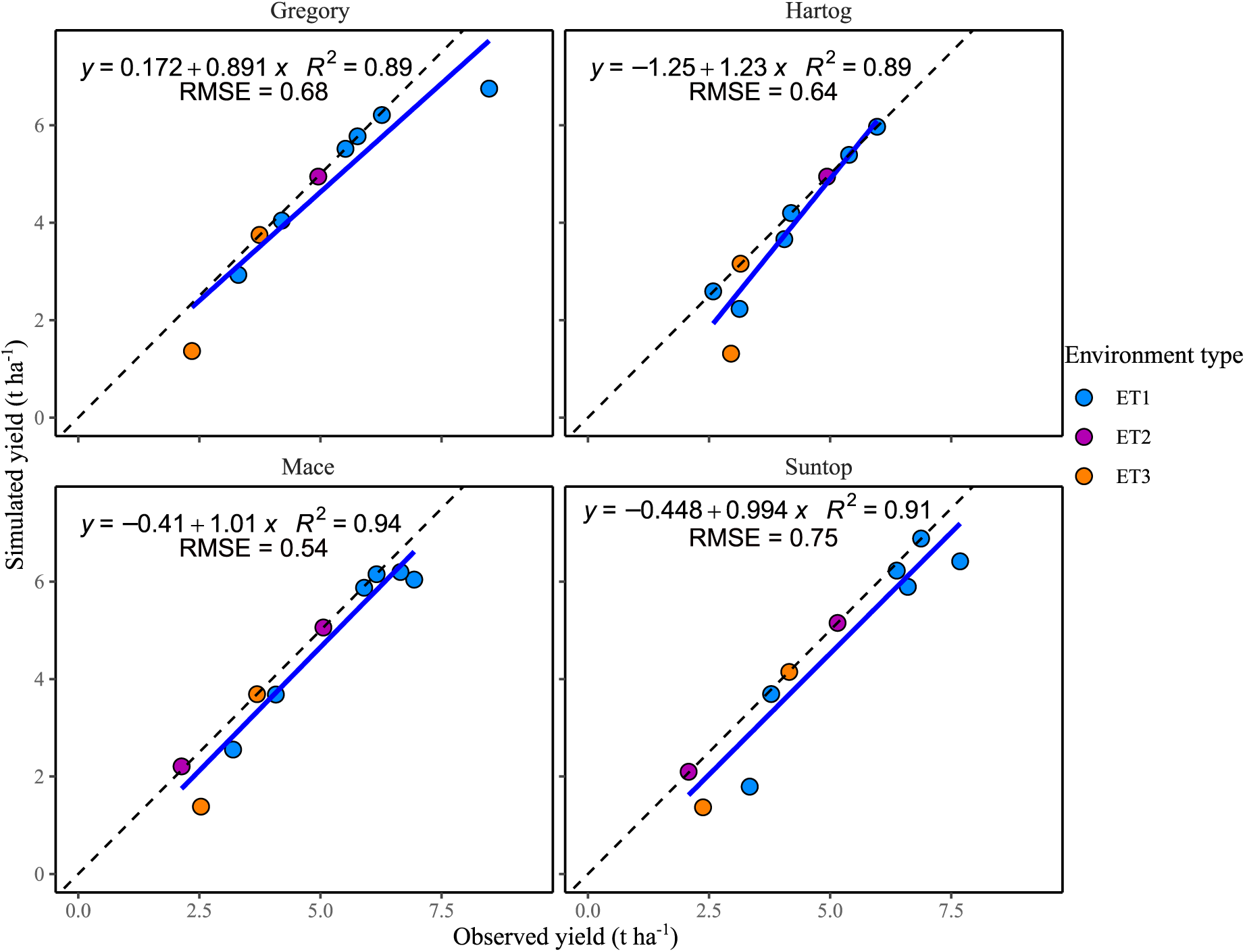
Simulated yield with the crop model against observed yield for four parental genotypes in the studied trials. To best simulate the genotype growth and development at each trial, the simulated flowering time was tuned to match observed flowering time, and soil moisture at sowing was optimized to decrease the deviation between observed and simulated yield. One unique soil moisture at sowing was used per site and year (i.e. same soil characteristics for all genotypes and treatments). Colors correspond to the environment types (i.e. ET1-ET3). For each genotype, the fitted relationship is presented as a solid blue line, and the 1:1 line is presented as a dashed black line.

**Fig. 4.**
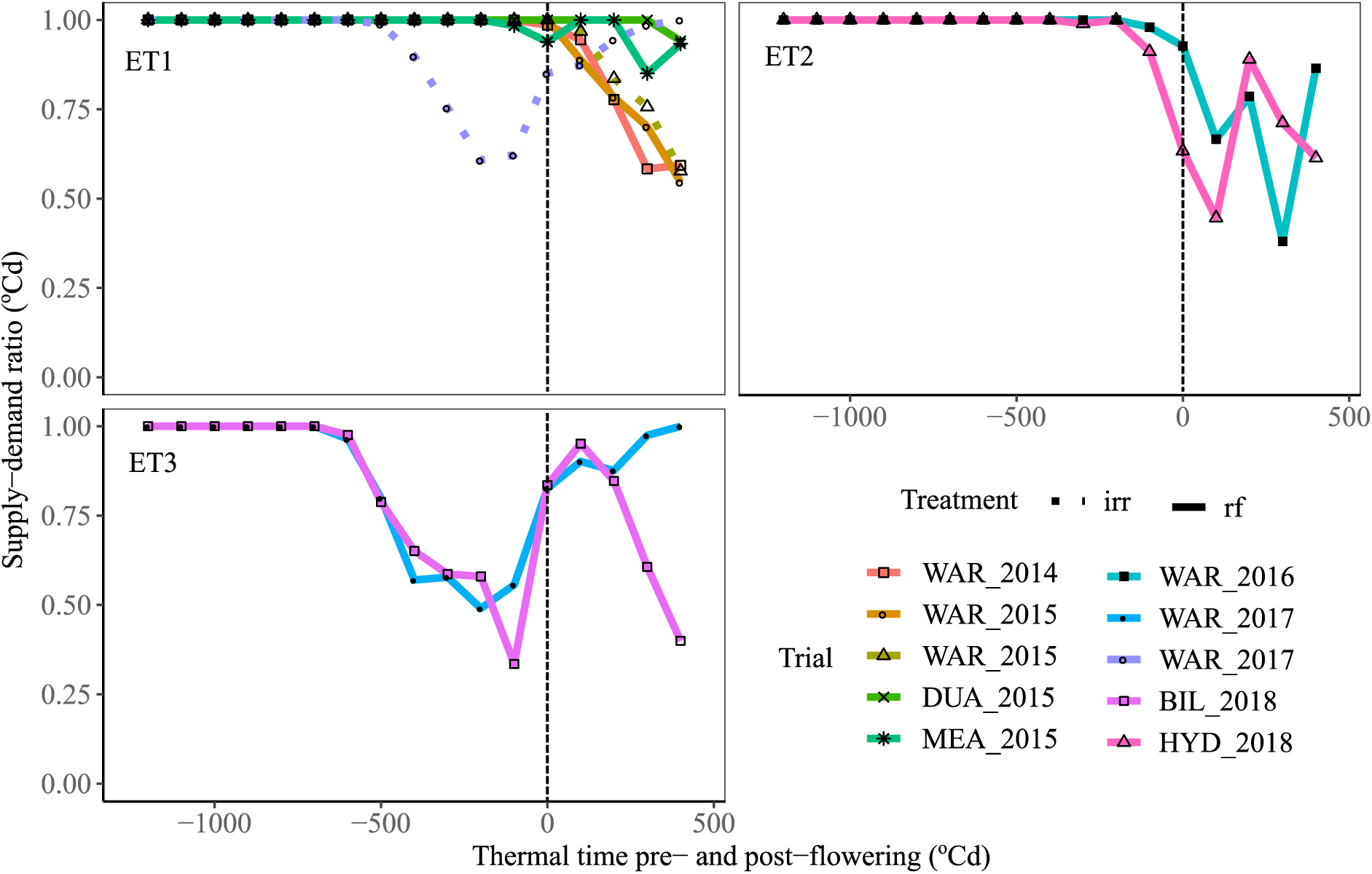
Water deficit patterns (supply-demand ratio) for Mace in studied trials classified by environment types (ET) as defined by Chenu et al. (2013). ET1 (no or minor water deficit), ET2 (post-flowering water deficit), and ET3 (water deficit starting pre-flowering and relieved during the grain filling). Line-type correspond to irrigated (irr, solid line) or rainfed (rf, dashed line) treatments.

### 2.6 Multi-environment trial analysis

A one step MET analysis was performed for yield using a linear mixed model, firstly without accounting for ET (Models 1 to 2, Table 3), and then with terms included to account for ETs (Models 3 to 4). Trial was fitted as a fixed effect, while the interaction of genotype and trial along with terms describing the structure of the experimental design for each trial were included as random effects. The spatial variation within each trial was modelled following the procedure of Gilmour et al. (1997). The analysis considered a series of *t* trials in which a total of *m* genotypes were evaluated. As such, 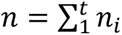 represent the total number of plots in the MET dataset, where *n*_*i*_ corresponds to the number of plots in trial *i* (*i* = 1, … *t*).

Specifically, the general form of Model 1 and 2, can be presented as

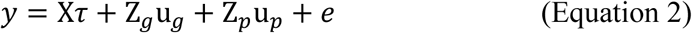

**Table 3.** Comparison of four different genotypic variance models fitted using the 10 trials and different variance structure applied to the G×E interaction effects (var(u_g_)). Likelihood and Akaike information criterion (AIC) of each model are presented together with N, which stands for number of parameters or, when indicated by an asterix (*), the number of non-boundary variance components. The model presented in bold font (Model 4) was selected based on lowest AIC value

| Model | Structure of $\text{var}(\mathbf{u}_g)$ | N | Log-likelihood | AIC |
| --- | --- | --- | --- | --- |
| 1 | $E \times G$ | 49* | 360 | -622 |
| 2 | $\text{diag}(E) \times G$ | 52* | 446 | -789 |
| 3 | $\text{diag}(ET) \times G + \text{diag}(E) \times \text{GRMp}$ | 62 | 734 | -1343 |
| 4 | **$\text{diag}(ET) \times G + \text{FA1}(E) \times \text{GRMp}$** | **77** | **914** | **-1673** |

Abbreviations: E, environment; G, genotype; diag, diagonal matrix; GRMp, genomic relationship matrix pedigree-based; $\text{var}(\mathbf{u}_g)$ , total variance; FA, factor analytic.
\* Number of non-boundary variance components.

where *y* is an *n* × 1 vector of trait values, *τ* is a vector of fixed effects (trial means), *u*_*g*_ is an *mt* × 1 vector of random effects for G×E interaction associated with design matrix Z_*g*_, *u*_*p*_ is the vector of random effects accounting for the experimental design structure in each trial, and *e* is the *n* × 1 vector of plot error effects.

The difference between Models 1 and 2 concerns the variance structure applied to the G×E interaction effects. In the case of Model 1, a simple variance component structure was utilised, whereby 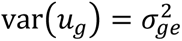. Alternatively, a diagonal variance structure was considered for the G×E interaction effects in Model 2, allowing for the estimation of heterogeneous genotypic variance for each trial 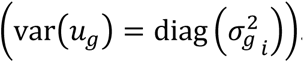.

To explore the value of envirotyping in terms of explaining some extent of the G×E interaction, an additional random term was added to the model to account for G×ET interaction. Assuming there are *q* ETs, a general form of Model 3 can be stated as

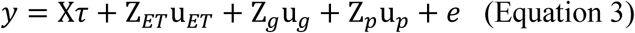

where *u*_*ET*_ denotes an *mq* × 1 vector of random genotype effects for each ET, with associated design matrix Z_*ET*_. All other terms are as defined in Equation 2. In this case, a diagonal variance structure was considered for the G×ET interaction effects in Model 3, allowing for the estimation of heterogeneous genotypic variance for each ET 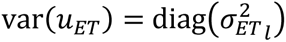, where *l* = 1, …, *q*.

As not all *m* genotypes were tested in each trial, it was necessary to have an adequate linkage between trials to estimate the genotypic covariance. Furthermore, we included the genetic relationship matrix based on pedigree information (GRMp) to enable an exploration of the additive and non-additive effects. Genotypic effects were partitioned using the genetic model of Oakey et al. (2006), whereby the genotypic effect was partitioned into additive (*u*_*a*_) using pedigree information, and non-additive components (*u*_*b*_). The vector of random G×E effects (*u*_*g*_) can then be written as

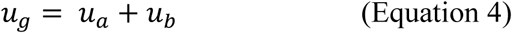

To this point, the variance structures considered have assumed trials to be independent. In order to allow for the estimation of genotypic covariance between trials, the factor analytic (FA) variance model (Smith et al., 2001; Kelly et al., 2009) based on *k* factors was used (Model 4, Table 3). Higher order FA models (i.e. with *k* value) were subsequently fitted to the G×E interaction effects until the most parsimonious model was reached, according to the Akaike information criterion (AIC).

The flexible FA model enabled the estimation of heterogeneous genotypic variance for each trial, along with heterogenous genotypic covariance between trials, via the representation of the G×E interaction effects as the product of environmental loadings and genotypic scores (Meyer et al., 2009).

Multi-environment trial analyses were performed using the ASReml-R software package (Gilmour et al., 2009) in the R statistical computing environment (R Core Team, 2018).

### 2.7 Genotypic correlation and genotype by environment (G×E) interactions

FA model (Model 4) was used to produce genotypic VCOV and correlation matrix to explain G×E interactions as a mode of heterogeneity in the breeding trials. The genotypic correlation matrix indicated the correlation between trials, which reflected the level of G×E; if the genotypic correlation between trials is high then G×E interactions are low and *vice versa*. In the FA model, factor loading increased the confidence in the data by stratifying the genotypes or environments such that there should be less G×E interaction within the ETs than between the ETs (Lin and Thompson, 1975). This assessment approach depends on the magnitude of G×E interaction, which indicates the performance of genotypes across environments, where the ‘environment’ corresponds to the sum of non-genetic factors that influence the phenotypic value associated with genotype. The factor ‘genotype’ refers to individuals from the families and founders of the MR-NAM populations. *e* is the vector of residual errors, representing unobserved or unmeasured factors that contribute to the variability in the response variable.

### 2.8 Trait and multi-trait analysis

A multi-trait analysis was performed for each of the five trials where stay-green measurements were collected, extending the approach of Ganesalingam et al. (2013). In a multi-trait analysis for each trial, the traits were considered as a fixed effect. The interaction between genotype and trait along with terms describing the structure of the experimental design, were included as random effects. All spatial trend, design and residual effects were modelled separately for each trait (i.e. flowering time, each of the stay-green traits, and yield). A FA model was used to estimate the genotype by trait effects, beginning with a model of order one. Higher order FA models were subsequently fitted to the genotype by trait effects until the most parsimonious model was reached, according to a log-likelihood ratio test. The FA model allowed heterogeneity of genotypic variance for each trait along with heterogeneity of genotypic correlations between all traits.

Heritability of yield was calculated by Equation 5 from the final Model 4:

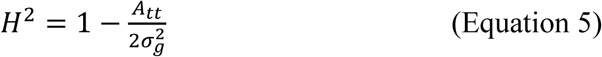

where *A*_*tt*_ is the average pairwise prediction error variance between genotypes for yield, and 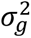 is the genotypic variance (Cullis et al., 2006).

Variance parameters were estimated using residual maximum likelihood (REML) estimation (Patterson and Thompson, 1971).

Predictions of genotype performance (yield) at each trial were generated from Model 4 as empirical Best Linear Unbiased Predictions (eBLUPs).

All multi-environment analyses were performed using ASReml-R version 3 (Gilmour et al., 2009), and all multi-trait analyses were performed using ASReml-R version 4 in the R environment (R Core Team, 2018).

A principal component analysis (PCA) was performed for each trial where stay-green traits were assessed to examine genotype by trait interaction between more than two traits.

Correlation between yield and each stay-green trait were estimated for each trial, and also averaged across trials within each ET for all genotypes. In addition, these correlations were calculated when only focusing on genotypes with similar phenology (similar number of days to flowering) in each trial. For each trial, these subsets of genotypes were chosen to have a flowering date close to the median flowering date of the trial and were denominated as the ‘sets selected for a restricted range of days to flowering’ (SRDF, Table 1).

The yield advantage associated with each of the stay-green traits was estimated as the average yield difference between genotypes with the 5% highest and genotypes with the 5% lowest value of the respective stay-green traits.

## 3 Results

### 3.1 Trial environments impacted yield and stay-green traits

Trial mean yield varied considerably across years and locations, with the mean yield ranging from 2.0 t ha^-1^ in HYD18rf to 7.2 t ha^-1^ in WAR14rf (Table 1). For a single location (Warwick, Queensland), the inter-annual range in average yield for rain-fed conditions varied from 3.8 to 7.2 t ha^-1^, i.e. a difference of 3.4 t ha^-1^ between the most extreme years.

The frequency of occurrence of drought ETs varied across trials and regions, with 60, 20, and 20% of the trials classified as ET1 (no or light water-deficit), ET2 (mild post-flowering deficit), and ET3 (severe water-deficit starting during the vegetative period and relieved during grain filling), respectively (Fig. 4). ET1 environments include two partially irrigated trials and four rainfed trials, while all ET2 and ET3 trials were rainfed. Most of the trials characterized as ET1 had deep soil types (Duaringa, QLD; Warwick, QLD) with more soil plant available water at sowing than other sites in most seasons (PAW, Table 2). Two trials were moderately water limited and characterized as ET2 (WAR16rf and HYD18rf). WAR16rf had the highest solar radiation values among all the trials, while HYD18rf had lowest PAW. Two severe stress environments (ET3) were observed in (i) WAR17rf with relatively dry soil conditions at sowing (PAW) and relatively low in-crop rainfall (Rain, Table 2), and (ii) BIL18rf with shallower soil, the lowest plant available water capacity (PAWC) (69 mm) among all the trials, and low in-crop rainfall (Rain, Table 2). While trials with more in-season rainfall tended to have less water-deficit (ET1 - ET2), rainfall was not the only key factor driving the water-deficit pattern. For example, the BIL18rf trial (Southern-east Queensland) received 105 mm in-crop rainfall and was severely water limited (ET3), while DUA15rf (QLD) only received 55 mm rainfall and was characterized as ET1, in part due to a higher PAW at sowing (165 mm) than BIL18rf (69 mm) (Table 2).

Average yield tended to decrease with the severity of water stress (from ET1 to ET3), while no clear trend was observed for flowering time (Table 1 and Fig. S2). Senescence occurred more rapidly in environments with severe water stress starting during the vegetative period (ET3) than in mild terminal stress (ET2) and water-sufficient environments (ET1), resulting in shorter periods from flowering to onset (OnS), mid (MidS) and end (EndS) of senescence in ET3 compared to ET1 and 2 (Fig. S2). No clear trend was observed for the initial value of NDVI which indicated the maximum value of greenness (Nmax).

### 3.2 Environmental characterization helped to untangle Genotype × Environment interactions and increased genotypic variance within environment types

The large range of environments tested resulted in high G×E interactions, and the genotypic correlation for yield between trials ranged from 0.10 to 0.81 (Fig. 5). Low genotypic correlations were found even between some years within the same site (Warwick, QLD). The broad-sense heritability for yield (*H*^2^) was variable between trials, ranging from 0.20 to 0.51 (Table 4).

**Fig. 5.**
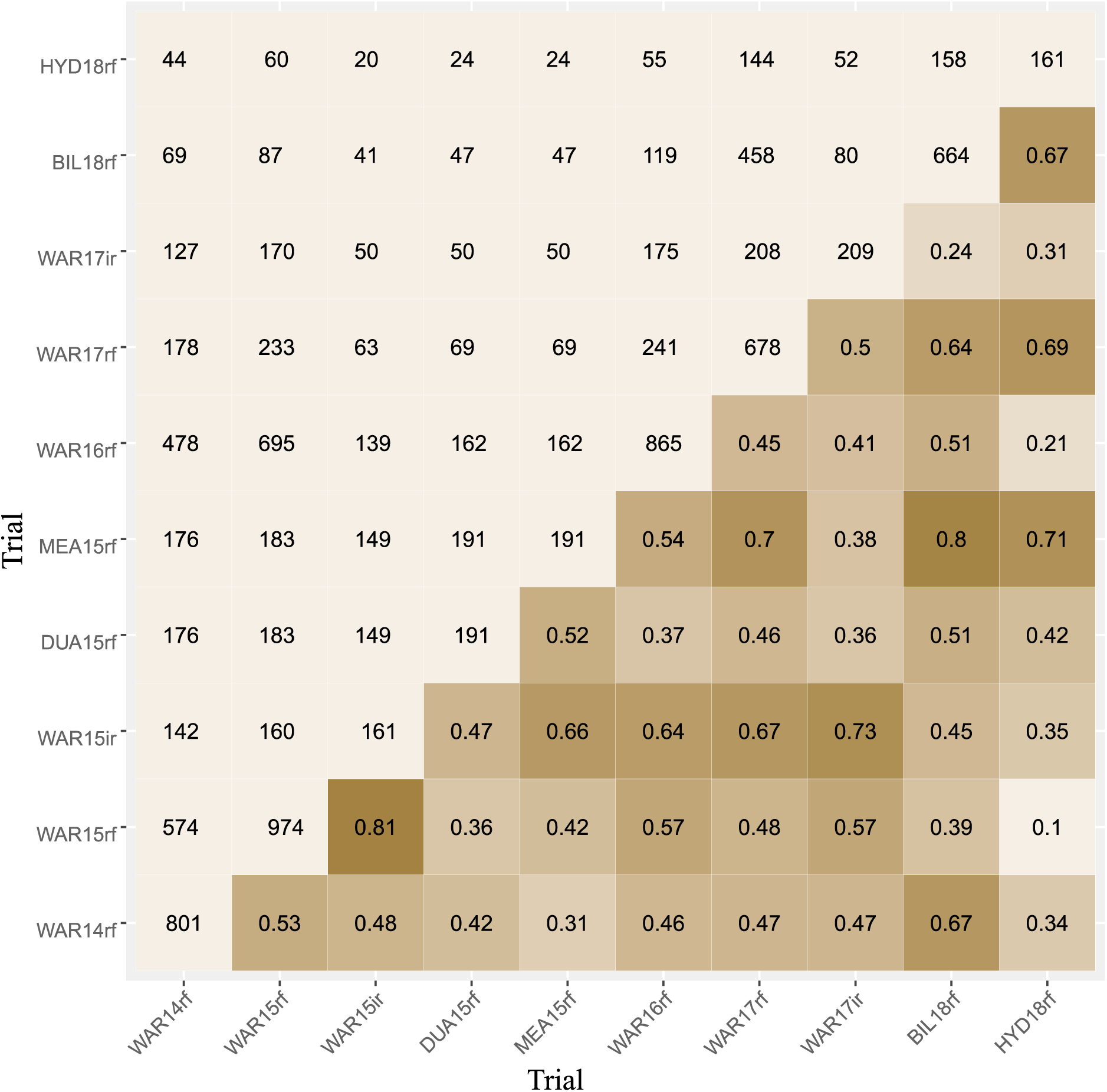
Heatmap for genotypic correlations from Model 2 (Table 3) for yield and wheat genotypes evaluated in the 10 studied trials. Upper diagonal indicates the number of shared genotypes evaluated in each pair of trials, while lower diagonal indicates genotypic correlations among pairs of trials. Trial characteristics are given in Tables 1 and 2.

**Table 4.** Broad-sense heritability (H^2^) for yield in each of the 10 trials.

| **Trial** | **$H^2$** |
| --- | --- |
| WAR14rf | 0.34 |
| WAR15rf | 0.48 |
| WAR15ir | 0.51 |
| DUA15rf | 0.20 |
| MEA15rf | 0.33 |
| WAR17ir | 0.30 |
| WAR16rf | 0.41 |
| HYD18rf | 0.23 |
| WAR17rf | 0.40 |
| BIL18rf | 0.34 |

After including ET information in the FA model of rank 1 (Model 4, Table 3), genotypic correlations between trials were generally greater within ET1 (0.27 to 0.78), ET2 (0.28), and ET3 (0.63). The higher genotypic correlations within ET indicate low G×E interaction compared to the correlation with other ETs (Fig. 6b). A Wald statistics test indicated a significant (p < 0.001) effect of genotype by ET interactions for yield, despite the fact that ET1 was highly correlated with ET2 (0.90) and ET3 (0.77; Fig. 6a). A more moderate correlation (0.45) was found between post-flowering stress ET2 with pre-flowering stress ET3 (Fig. 6a). This may be due to the different nature of the water-deficit stress, which may have involved different adaptative mechanisms influencing genotypic variation for yield.

**Fig. 6.**
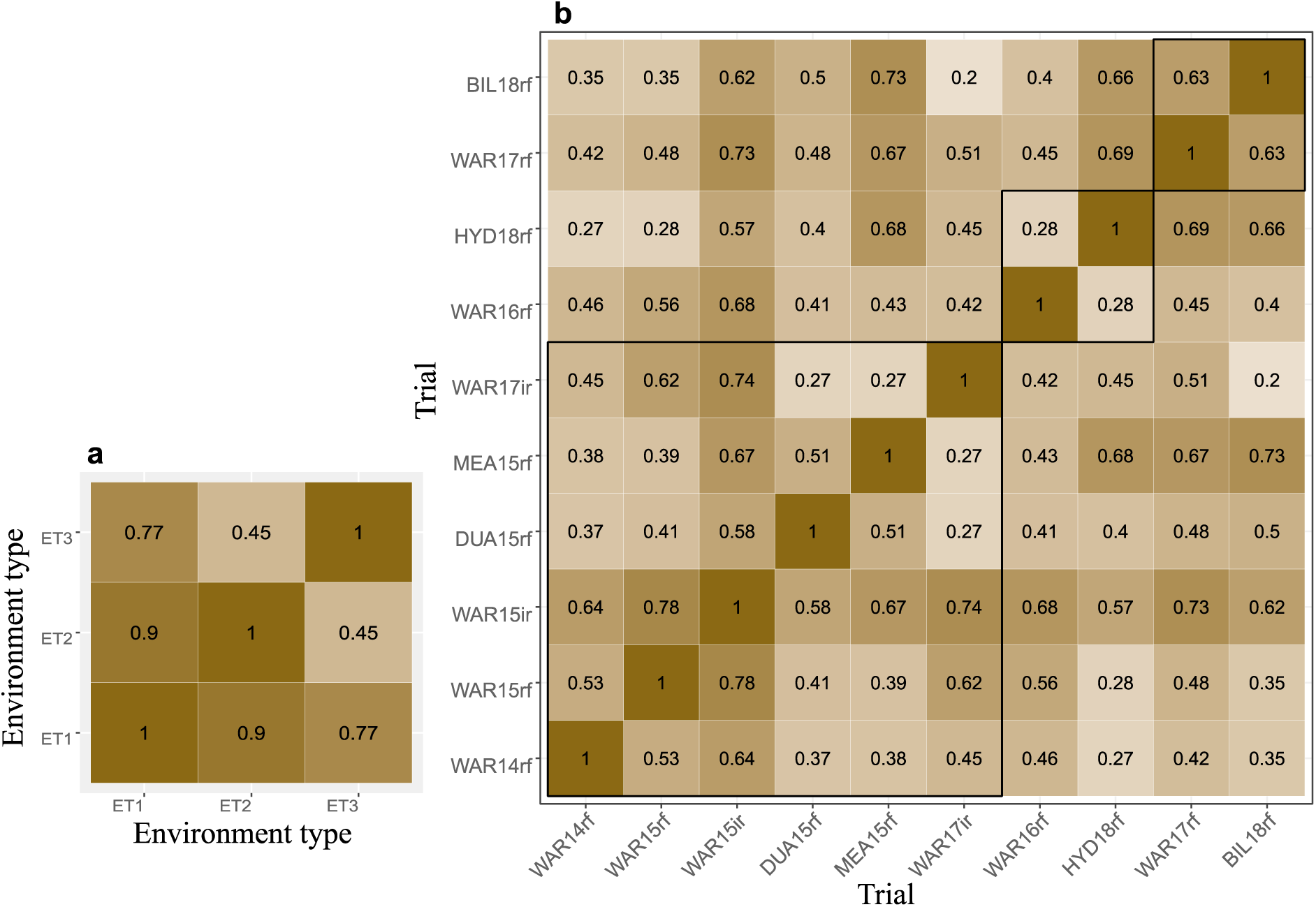
Heatmaps of the genotypic correlations estimated from Model 4 (Table 3) for yield across (a) environment types (ET) and (b) trials. Trials are ordered by environment types (ET1 (left) to ET3 (right)). In (b), black bold lines encircle trials within each of the ETs.

By including the ET information in the linear mixed model (Model 4 vs Model 2; Table 3), estimates of genotypic variance for yield increased by 41% to 87% depending on the trial (Fig. 7c). On average, genotypic variances increased by 66% in ET1, 67% in ET2 and 75% in ET3 (Fig. 7d). Better estimation of genotypic variance provides a basis for increases in the prediction accuracy.

**Fig. 7.**
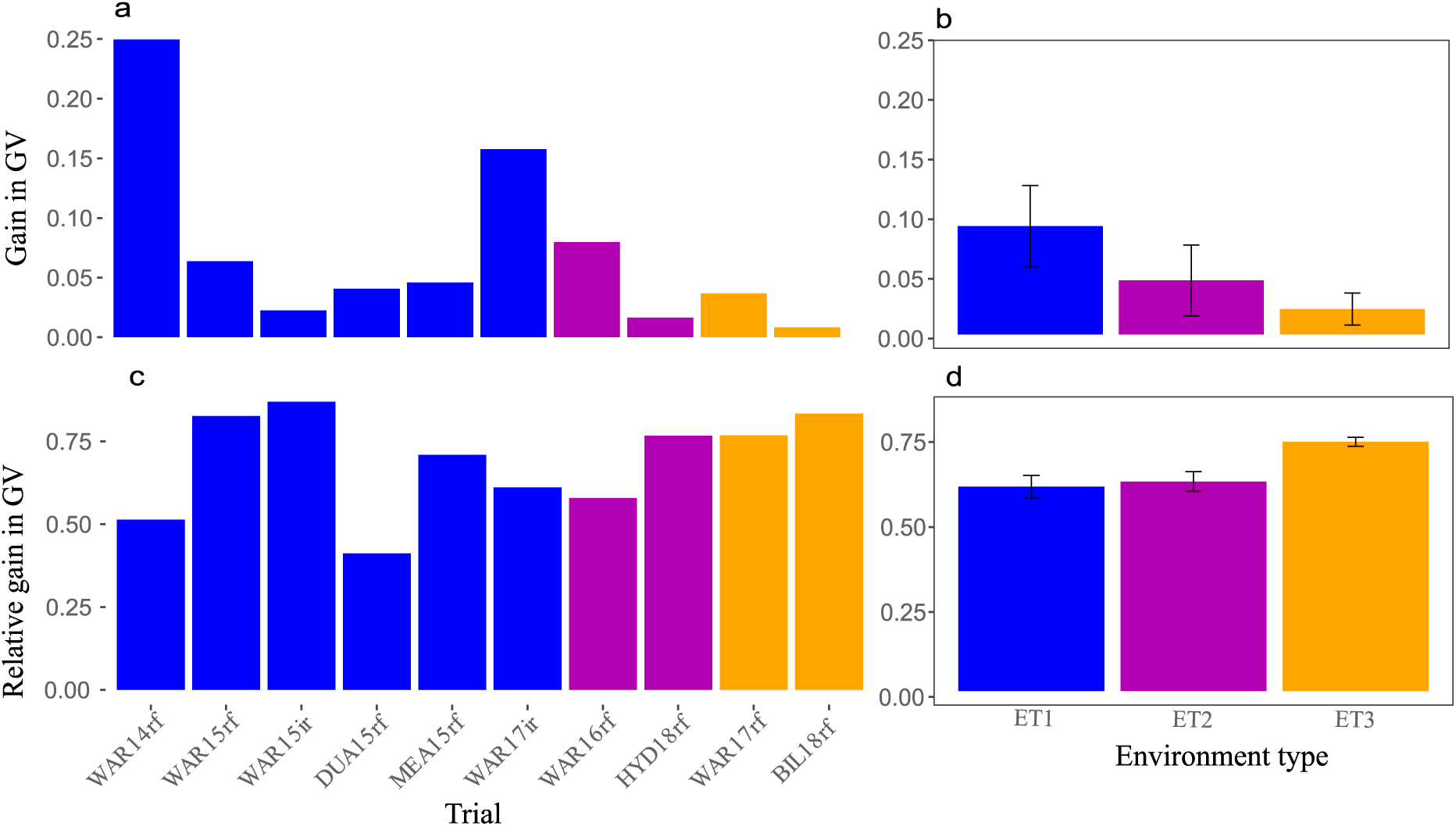
Gain in genotypic variance (GV) for yield when adding envirotyping information to the statistical model (i.e. Model 4 vs Model 2, Table 3) and presented in absolute value (a-b) and in relative value (ratio, c-d) by (a, c) trials and (b, d) environment types, for the 10 trials. (a-b) The gain in genotypic variance was estimated as the difference between genotypic variance with ET (Model 4) and without ET information (Model 2; Table 3). (c-d) The relative gain in genotypic variance was estimated as the ratio between the gain in genotypic variance and the genotypic variance without ET information (Model 2). Error bars correspond to standard error.

### 3.3 Response of genotypes for stay-green traits in different environment types

Principal components analysis was used to investigate the relationship between stay-green traits, flowering, and yield within trials (Fig. 8). The first two principal components (PC) combined explained more than 70% of the total variance in all studied environments. PC1 explained over 50% of the variance in all cases while PC2 explained substantially less (Fig. 8). Eigenvectors in the graphs showed the correlation value across yield, stay-green traits, and flowering for each trial. In most of the trials, yield and flowering were negatively correlated with each other, except WAR15rf (ET1). Overall, positive correlations were observed between stay-green traits and yield, with a high correlation for stay-green traits OnS (onset of senescence), MidS (mid senescence), and to a lesser extend SGint (stay-green integral; Fig. 8). Vectors representing OnS, MidS, and SGint were largely aligned with the yield, but the length of the stay-green trait vectors was greater compared to the yield vector (Fig. 8).

**Fig. 8.**
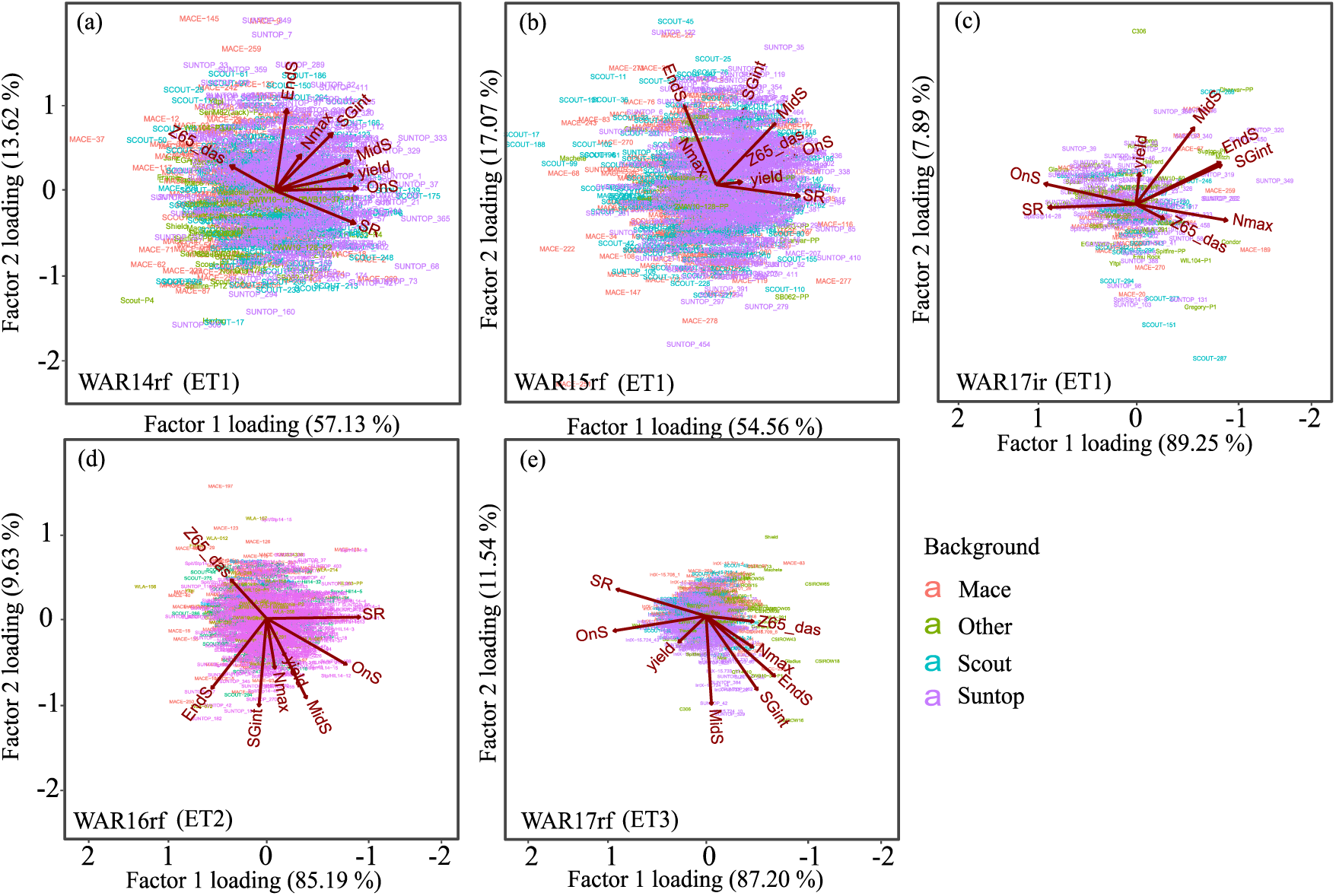
Biplots for the seven trials where stay-green was measured. The biplots summarise correlations between stay-greens traits (OnS, MidS, EndS, SR, Nmax, and SGint), flowering time in days after sowing (Z65_das) and yield. Vectors indicate the traits. Small angles between vectors indicate positively correlated traits. Points indicate genotypes. Color corresponds to the different genotypic backgrounds to which the genotypes belong (e.g. Mace, Scout, and Suntop).

For ET1 environments, two out of three tested trials, vectors for SR (indicator of senescence rate) were in a similar direction to the yield vector, but not as close as OnS, MidS, and SGint vectors, which indicates a weaker correlation with yield. A large number of genotypes that clustered at the centre of the biplot exhibited relatively homogenous characteristics, while genotypes towards the extremes of the biplots had dissimilar characteristics. Stay-green and yield vectors were close to each other, indicating a positive correlation in water-sufficient environments (ET1) with the exception of some stay-green traits in WAR17ir. This agrees with the significant and relatively high positive genotypic correlations observed between stay-green traits (except EndS) and yield in ET1 (Fig. 9).

**Fig. 9.**
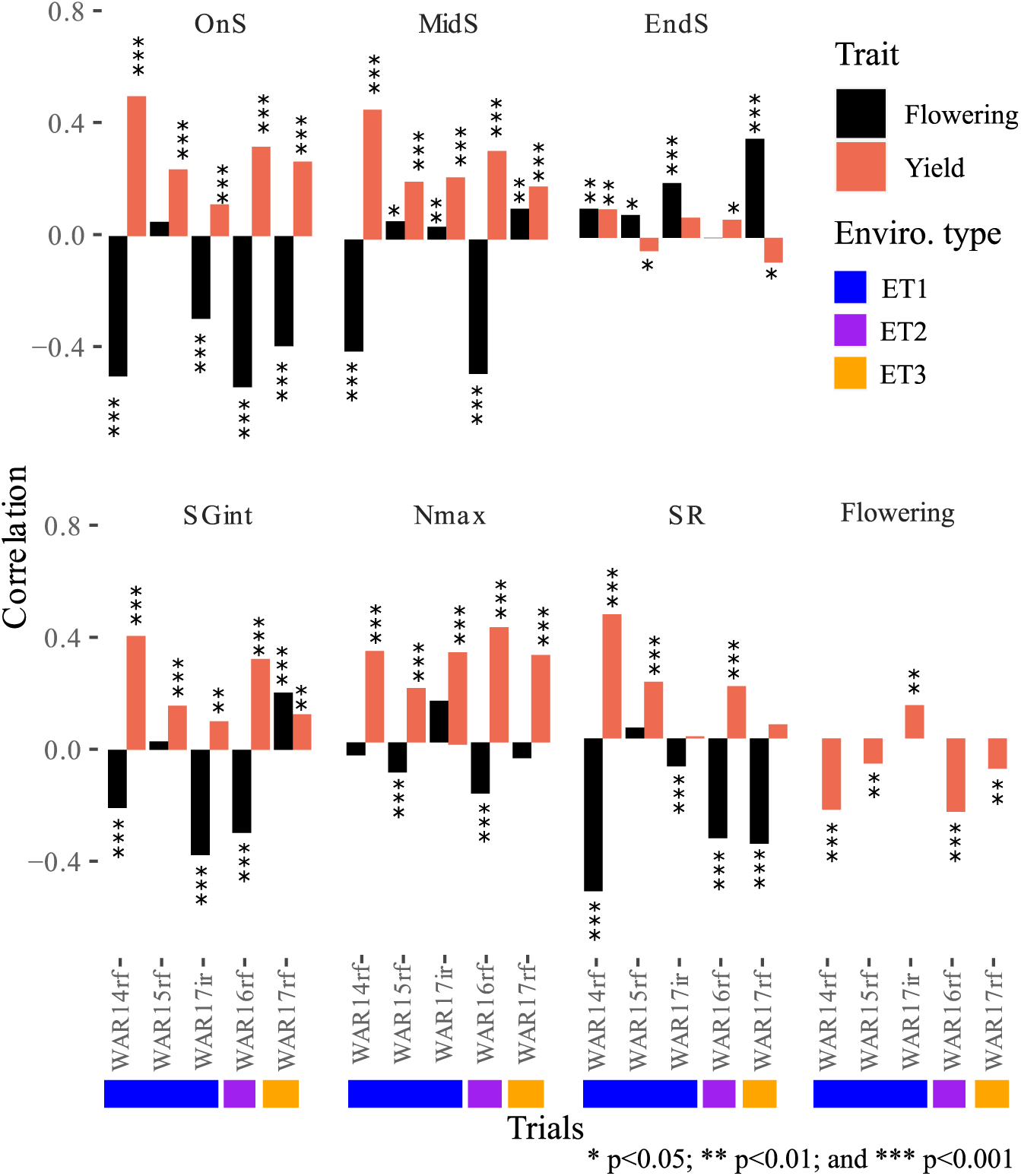
Correlations between stay-green traits with flowering (black bars) and yield (coral bars), and correlations between flowering and yield (coral bars) in different trials. Trials are ordered in ascending order of water stress, i.e. ET1 (blue), ET2 (mauve), and ET3 (orange). Stay-green traits represented are; start of senescence (OnS), mid senescence (MidS), end of senescence (EndS), indicator of the rate of senescence (SR), maximum greenness (Nmax), and stay-green integral (SGint). ***, **, and * indicate the 0.001, 0.01, and 0.05 level of significant difference, respectively.

In ET2 (WAR16rf), post-flowering water-stress was more severe than in ET1 (Fig. 4). A strong positive relationship was observed between MidS and yield, as the vectors of MidS and yield had similar directions, but the yield vector was smaller in length. OnS, MidS, and SGint were also positively correlated to yield.

In ET3 (WAR17rf), genotypes experienced severe pre-flowering stress relieved after flowering (Fig. 4). Genotypes with higher values of onset of senescence (OnS) correlated with lower values for days to flowering (Z65_das), which suggests early flowering genotypes started senescing later, possibly to due the fact that the stress occurred at a later stage for those genotypes. Vectors for OnS and MidS were closer to that of yield compared to other stay-green traits (Fig. 8), indicating stronger positive correlations of OnS and MidS with yield. Nmax, SGint and EndS had a positive correlation with flowering which suggests that late flowering genotypes had higher values for Nmax, SGint, and EndS.

Overall, results indicated that yield closely related to MidS - OnS > SGint Nmax in most of the environments (Figs. 8-9).

### 3.4 Influence of flowering time on stay-green traits and their correlation with ***yield***

Genotypes substantially differed in phenology, with duration from sowing to flowering time varying by 15-27 days (WAR17ir - WAR15rf) between the earliest and latest genotypes (ORDF, Table 1). Flowering time significantly correlated with some stay-green traits and yield in all ETs (Fig. 9). In most trials, late-flowering genotypes tended to start senescing earlier (OnS) but with a lower rate of senescence (SR) so that they tended to end senescing later (EndS) and with lower yield than early-flowering genotypes. This could be partly due to the fact that those late-flowering genotypes were affected by water stress at an earlier developmental stage than early-flowering genotypes, as all studied trials were subjected to water limitation (with different intensity and timing; Fig. 4).

Focusing on genotypes with similar phenology (i.e. which experienced the stress at a similar developmental stage) by selecting genotypes with a narrower range of value for days to flowering (SRDF) can help to estimate correlations between stay-green traits and yield with a reduced confounding effect due to flowering date (Fig. 10). For genotypes with similar flowering time within each trial, no significant correlations were observed between flowering and yield (except in WAR14rf). However, there were significant correlations found between stay-green traits and yield in most trials for the same genotypes (Fig. 10). Importantly, considering only genotypes with similar phenology (SDRF) significantly increased correlation values between stay-green traits (OnS, MidS, SR, EndS and SGint) with yield in all ETs, (Figs 9-10). Overall, for genotypes with similar phenology, stay-green traits were significantly and positively correlated with yield in all ETs (Fig. 10). The strength of the correlations among the ETs had a tendency to be stronger in ET1 > ET2 > ET3 (Fig. 10).

**Fig. 10.**
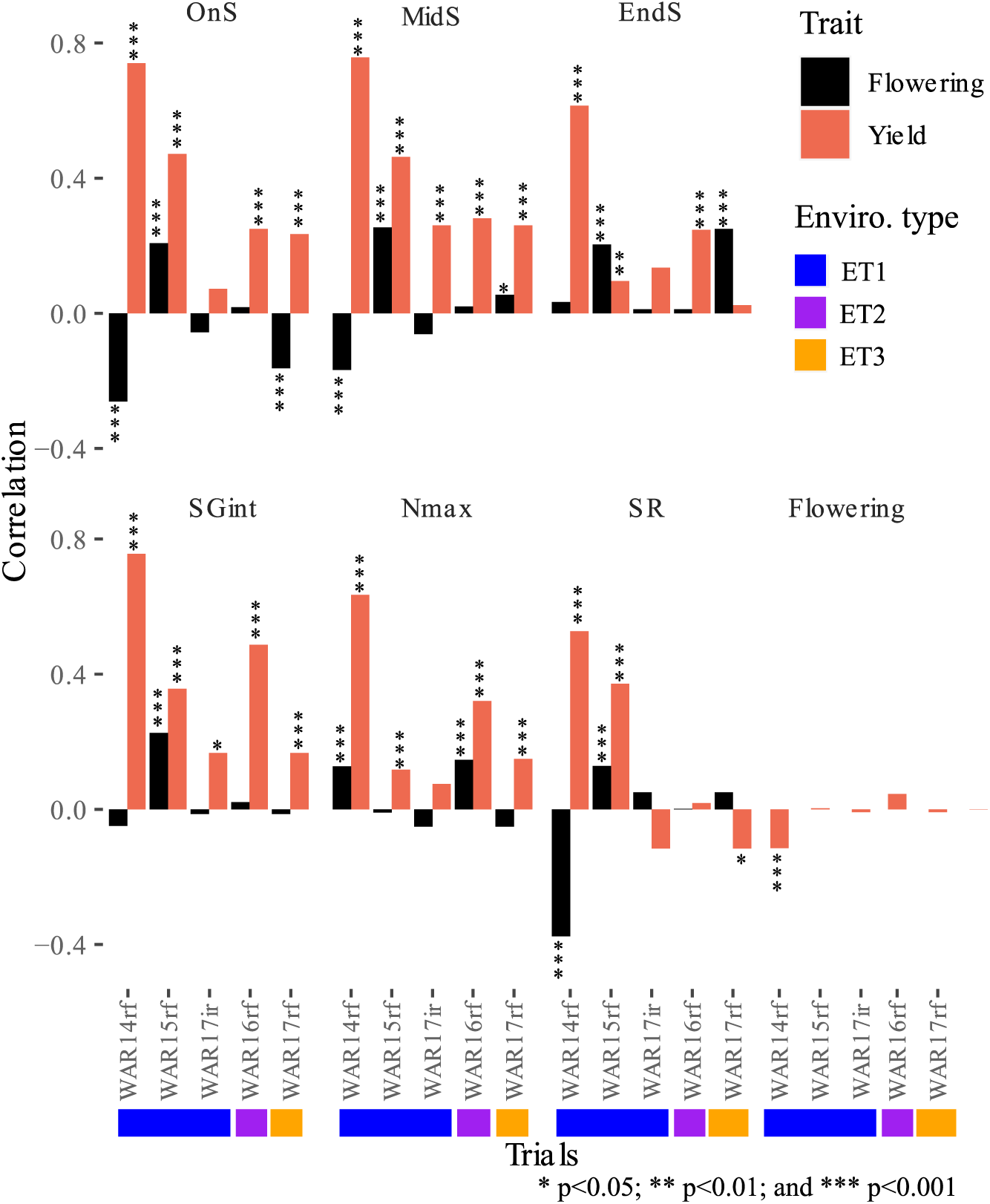
Correlations between stay-green traits with flowering (black bars) and yield (coral bars), and between flowering with yield (coral bars) for subsets of genotypes with similar duration to flowering (SRDF, Table 1) selected from each trial. Trials are ordered in ascending order of water stress, i.e. ET1 (blue), ET2 (mauve), and ET3 (orange). Stay-green traits represented are; start of senescence (OnS), mid senescence (MidS), end of senescence (EndS), indicator of the rate of senescence (SR), maximum greenness (Nmax), and stay-green integral (SGint). ***, **, and * indicate the 0.001, 0.01, and 0.05 levels of significant difference, respectively.

### 3.5 Yield advantage of stay-green traits

Yield advantage was estimated for each stay-green trait by calculating the average yield difference between genotypes having the 5% lowest and 5% highest values for the considered stay-green trait (Fig. 11). In all ETs, stay-green traits contributed positively to yield. Stay-green traits (excluding EndS) contributed an average yield advantage between 0.7 to 1.1 t ha^-1^ in ET1 (9.7 to 15.9%), slightly less in ET2 (0.42 to 0.81 t ha^-1^, i.e. 8.7 to 16.6%) and in ET3 (0.23 to 0.73 t ha^-1^, i.e. 6.0 to 18.9%).

**Fig. 11.**
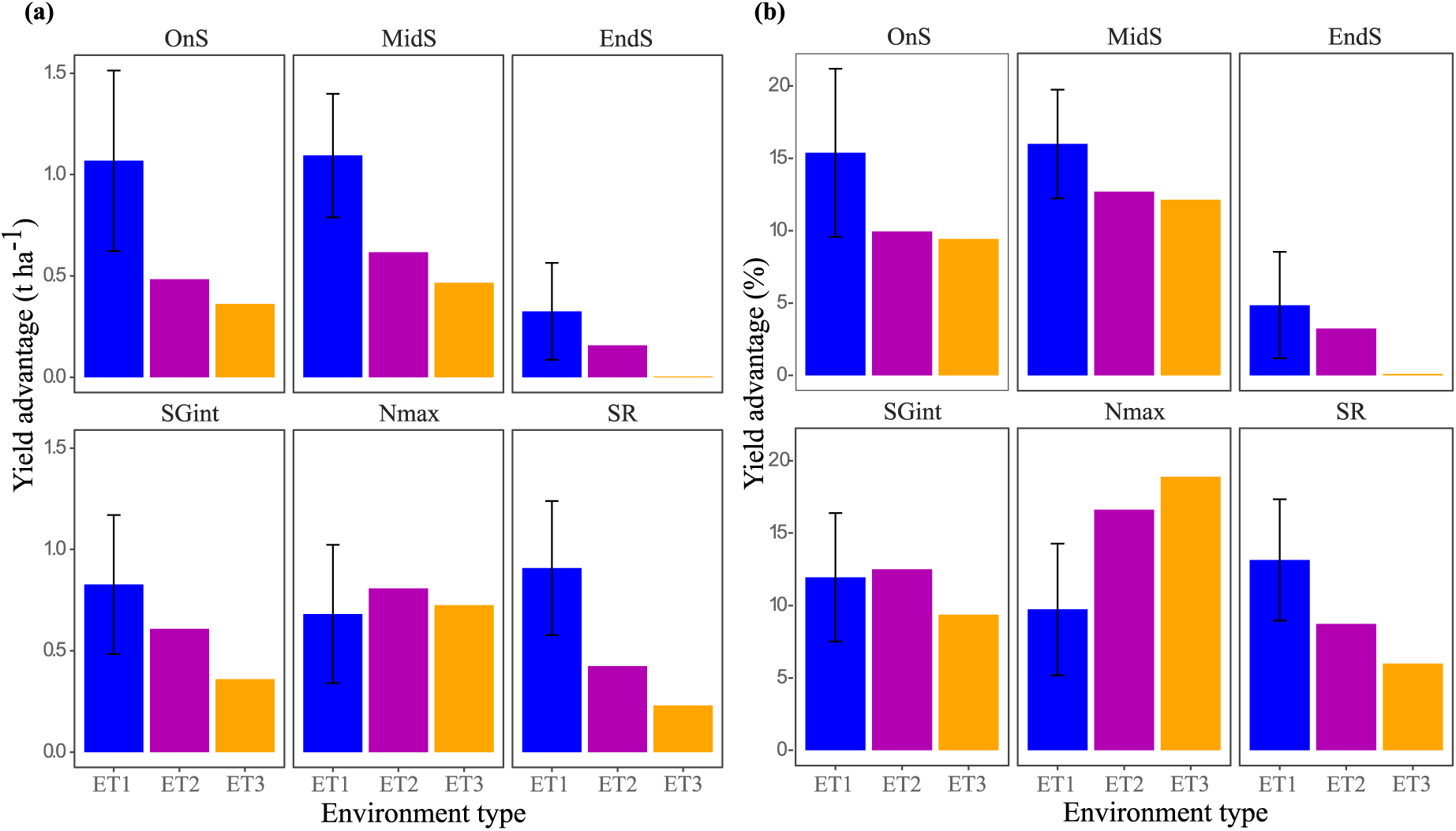
Yield advantage associated with the stay-green traits in the different environment types and presented in (a) absolute value (t ha^-1^) or (b) relative value (%). Yield advantage was estimated as the yield difference between genotypes with the 5 % highest and the 5% lowest value of the respective stay-green traits (i.e. tails of the population). Stay-green traits are start of senescence (OnS), mid of senescence (MidS), end of senescence (EndS), indicator of the rate of senescence (SR), maximum greenness (Nmax), and stay-green integral (SGint). Color corresponds to the environment types (ETs). Bars correspond to standard error.

In ET1, OnS and MidS had highest yield advantage (1.09 and 1.06 t ha^-1^, respectively) followed by SR, SGint, and Nmax. In ET2, Nmax had higher yield advantage (0.81 t ha^-1^, 16.6%) followed by other stay-green traits (Fig. 11). In ET3, OnS and MidS were the closest stay-green traits associated with yield in the biplot (Fig. 8) and were associated with a yield advantage of 0.36 and 0.46 t ha^-1^ (9.4 and 12.1%), respectively (Fig. 11). Nmax was also highly correlated to yield in ET3 (Fig. 8) and a high yield advantage (0.73 t ha^-1^, 18.9%; Fig. 11). Overall, results indicate a substantial role of stay-green traits in yield realisation across ETs.

## 4 Discussion

### 4.1 Multi-environment trials with substantial G×E interactions

The studied wheat MR-NAM population was developed to investigate the impact of adaptive traits like stay-green and their underlying genetic controls in response to heat and drought (Richard, 2018). Here, 10 trials conducted under irrigated and rainfed conditions were studied in a multi-environment analysis to evaluate genotypic variance and G×E interactions for yield performance. While yield was likely affected by several stresses (Chenu et al., 2013; Ullah et al., 2024; Yahya et al., 2024), a major influence was the role of water deficit, which created different patterns in the timing and intensity of water stress during crop development (Fig. 4). The trial environments sampled were assigned to three of the 4 major ETs previously described for the Australian TPE (Chenu et al., 2013). Of the 10 trials, 60% (6) were classed as ET1 with smaller numbers (2) in post-flowering deficit environments ET2, and two trials in the more severe water-deficit ET3 with the deficit starting during the vegetative development. A high degree of variation for yield remained within each of these ETs (Fig. 4). Genotypic correlations were used to characterize the relative expression of G×E interactions, where low correlations indicate low stability, which suggested genotype rank in one environment was different than others. Genotypic correlation between the trials (Fig. 5) ranged from 0.10 (WAR15rf, HYD18rf) to 0.81 (WAR15rf, WAR15ir), indicating the high to low G×E interaction among trials, respectively. While estimation of genotypic covariance/correlation is also dependent on the number of genotypes in common between trials (Fig. 5), genotypic covariance could still be computed as trials shared enough common genotypes. Low genotypic correlation and variations in heritability in G×E analysis suggest a single ranking could not be used across all trials (Fig. 5; Table 4). Less than half of the genotypic correlations were considered relatively high (>0.50) between the trials.

### 4.2 Potential of environmental characterization to reduce the proportion of unexplained G×E interactions

Previous attempts have been made to incorporate climate and soil factors into linear mixed models (Vargas et al., 2007). However, these statistical models do not consider the interaction between the crop and the environment, for example the feedback of plant growth on soil water depletion (Chenu et al., 2011). Furthermore, selection based on yield without considering the environment could result in discarding genetic material that offers value in under-represented or specific ETs that are, nevertheless, important for the TPE.

To investigate the value of envirotyping to interpret G×E interaction for yield, ET information was included in the linear mixed model framework (Table 3). Including ET information in the model (Model 4 vs Model 2, Table 3) resulted in high gain in explained genotypic variance in most trials, with a 75% gain in the proportion of explained genotypic variance in ET3 environments experiencing severe stress and slightly smaller gains in ET2 and ET1 (Fig. 7d). Similarly, inclusion of ET information was reported to assist G×E untangling in other studies (e.g. in wheat, Chenu et al., 2011). It also helped to explain >30% of the genotypic variance in a sorghum MET (Chapman, 2008). Despite the substantial added value of incorporating ET into the model, a high degree of genotypic variation remained for trials within ETs. This may have been partly due to lower numbers of concurrent genotypes evaluated in some pairs of trials (Fig. 5) and was attenuated by fitting pedigree information (GRMp) into the linear mixed model.

Estimation of genotypic covariance (correlation) is dependent on the number of genotypes in common between trials. When trials have less genotypic connectivity as BIL18rf and HYD18rf with the other trials (Fig. 6), genotypic correlation may be computed but can be relatively unreliable (Dodds et al., 2019). This highlights the advantage of balanced trial designs for G×E analysis and plant breeding to ensure robust estimation of genotypic VCOV structure and reduce the risk of discarding valuable germplasm (Van Eeuwijk et al., 2001). However, such balance is often missing in historic datasets as it is often not a high priority for resource allocation in a commercial breeding program.

The highly variable and unpredictable nature of weather across the Australian wheatbelt makes it difficult to control or predict ET for rainfed yield trials (Zheng et al., 2018). For example, ET4 was not represented in the trials analysed in this study while ET1 was over-represented in comparison to the TPE (Chenu et al., 2011, 2013). To improve broad adaptation, germplasm should be evaluated in a range of environments, particularly for drought-prone target environments. A thorough and more balanced sampling of ETs can enable more thorough G×E analyses and provide breeders more options to achieve genetic gain. In a practical sense, application of envirotyping in a breeding program enables breeders to weight the performance of genotypes according to the trial growing environments compared to the TPE of the breeding program (Podlich et al. 1999; Chenu et al., 2011). Weighted selection strategies have been demonstrated to be particularly beneficial in target environments that experience a high degree of G×E interaction (Podlich et al. 1999, Messina et al. 2023).

Overall, inclusion of ET improved estimation of genetic performance and understanding of G×E, even in the studied unbalanced dataset. The method has considerable potential to improve estimation of genotypic variance for selection in breeding programs using unbalanced historical and on-going trials for highly variable climates.

### 4.3 Benefits from stay-green traits to enhance germplasm yield performance

Stay-green traits were characterized by fitting a logistic model to weekly NDVI measurements indicating canopy greenness for each plot of five trials with contrasting water-stress levels (Fig. 2; Christopher et al., 2014). All studied stay-green traits (except EndS) were positively correlated with yield in all ETs (Figs. 9-10). The strength of the correlations between the stay-green traits and yield varied with the patterns of water stress in the trials (Fig. 9). Focusing on a subset of genotypes with similar flowering time within each trial enhanced correlations between stay-green traits and yield (Fig. 10). Results thus indicated that the significant correlation of stay-green traits with yield was not likely derived from the correlation between flowering and yield (Fig. 10). This agreed with findings from a previous study using a biparental population in which variations for flowering dates was reduced by selection prior to experimentation (Christopher et al., 2016).

OnS, MidS, SGint, and Nmax were associated with yield in both water-sufficient and water-limited environments (Figs. 9-10). For instance, Nmax had a yield advantage consistently above 0.7 t ha^-1^ in ET1, ET2, and ET3 (Fig. 11). Hence, this single trait was associated with an average yield gain of 10% in water-sufficient environments ET1 and 16-18% in water stressed ETs, while yield breeding improvement of broadacre crops like wheat are rarely above 1-2% per annum (Duvick et al., 2004; Ray et al., 2014; Fisher et al., 2014; Messina et al., 2023). OnS and MidS were found more influential for adaptation to water-sufficient (ET1; 15% yield gain) than to water-limited environments (ET2 and ET3, ∼10% gain; Fig. 11). EndS was weakly correlated with yield in all ETs, which reflects that this trait, as it is calculated here, may not be as valuable for selection to improve adaptation in wheat (Fig. 9). The results agree with those for the wheat biparental population reported by Christopher et al. (2016). However, the results for wheat contrast with those found in sorghum (Borrell et al., 2014), a crop with a more perennial habit, and for which EndS was highly correlated with yield in post-flowering drought environments (ET3; Jordan et al., 2012).

In crops like wheat and maize, stay-green has been associated with genetic yield advance made over the past decades (Chenu et al., unpublished; Lee and Tollenaar, 2007). Stay-green appears to be the main photosynthetic trait associated with higher yield potential in wheat cultivated under good agronomic conditions (Carmo-Silva et al., 2017). In some dry environments, stay-green appears as a proxy or an emergent property for the multiple traits that enable different genotypes to avoid drought stress in crops like wheat or sorghum (Borrell et al., 2014). Stay-green has also been reported to have value on yield in response to heat stress (Lopes and Reynolds, 2012; Pinto et al., 2016; Ullah and Chenu, 2019; Yahya et al., 2022). However, a constitutive stay-green is not necessarily always beneficial, as reported in drought-prone Mediterranean environments (Chairi et al., 2020). Extending the duration of grain filling can work against the phenological adjustment (and thus a shorter crop cycle) that is typical of the genotypic adaptation of wheat to drought-prone environments (Loss and Siddique, 1994; Araus et al., 2002), and holistic Genotype × Environment × Management (G×E×M) approaches are important to maximize long-term yield (Zheng et al., 2012; Kirkegaard et al., 2014; Hammer et al., 2014; Chenu et al., 2017, 2018; Zheng et al., 2018; Collins and Chenu, 2021).

Overall, in the tested environments, OnS, MidS, SGint, and Nmax appeared as traits substantially beneficial to yield, as indicated by their positive and consistent correlation with yield in all three ETs (Figs. 8-10). Thus, for the MR-NAM population, selection of OnS, MidS, and Nmax could be very beneficial to improve adaptation across environments. Progress in high-throughput phenotyping (e.g. Chapman et al., 2014; Rebetzke et al., 2016; Potgieter et al., 2017 and 2021) should assist selection of genotypes with traits that help delay senescence to better tolerate heat and drought stress. Selecting for stay-green traits may aid improvement of tolerance to water and heat stress which are both likely to worsen in major wheat producing regions in future due to climate change (Lobell et al., 2015; Watson et al., 2017; Ababaei and Chenu, 2020; Collins and Chenu, 2021).

## 5 Conclusion

For conventional and molecular-level selection, understanding the biophysical conditions of the environments in which genotypes are evaluated is important to effectively assess their value in the TPE. In this study, envirotyping for levels of water deficit helped to better explain genotypic variation for yield in a wheat MR-NAM and reduce G×E interactions to improve the predictability of the genotype performance. In addition, stay-green phenotyping further facilitated the understanding of the genotypic behaviour in different environments. In the MR-NAM population, genotypes with later OnS and MidS, as well as other stay-green traits had greater yield in both water-sufficient and water-limited environments. With progress in high throughput field phenotyping, selection for stay-green traits could assist the development of a broad-range adaptation for water-sufficient and water-limited environments. Selection in breeding populations could be improved by including an envirotyping and phenotyping approach to better understand G×E interaction and crop physiology in order to increase predictability of genotype performance for current and future climates.

## Abbreviations

°Cd: degrees Celsius days;
BLUPs: best linear unbiased predictors;
EndS: end-senescence;
ET: environment type;
G×E: genotype by environment interaction;
ICARDA: International Centre for Agricultural Research in Dry Areas;
MidS: mid senescence;
MR-NAM: multi-reference parent nested association mapping;
Ma-NAM: Mace derived nested association mapping;
NAM: nested association mapping;
NDVI: normalised difference vegetation index;
Nmax: maximum greenness;
OnS: onset senescence;
QDAF: Queensland Government, Department of Agriculture and Fisheries;
RILs: recurrent inbred lines;
Sc-NAM: Scout derived nested association mapping;
SGint: stay-green integral;
SR: indicator of maximum senescence rate;
Su-NAM: Suntop derived nested association mapping.

## Acknowledgements

The authors wish to thank the field research teams at the University of Queensland, Gatton research farm, and QDPI Queensland, and particularly Mr Scott Diefenbach, for excellent technical assistance.

## Author attributions

AA: Formal Analysis, Visualization, Writing – original draft. JC: Conceptualization, Investigation, Methodology, Resources, Supervision, Writing – review & editing. CF: Formal Analysis, Writing – review & editing. MC: Investigation, Supervision, Writing – review & editing. BM: Data curation, Formal Analysis. BC and KF: Supervision. LH: Conceptualization, Resources, Supervision, Writing – review & editing. KC: Conceptualization, Investigation, Methodology, Resources, Supervision, Writing – review & editing.

## Supplementary figures

**Fig. S1.**
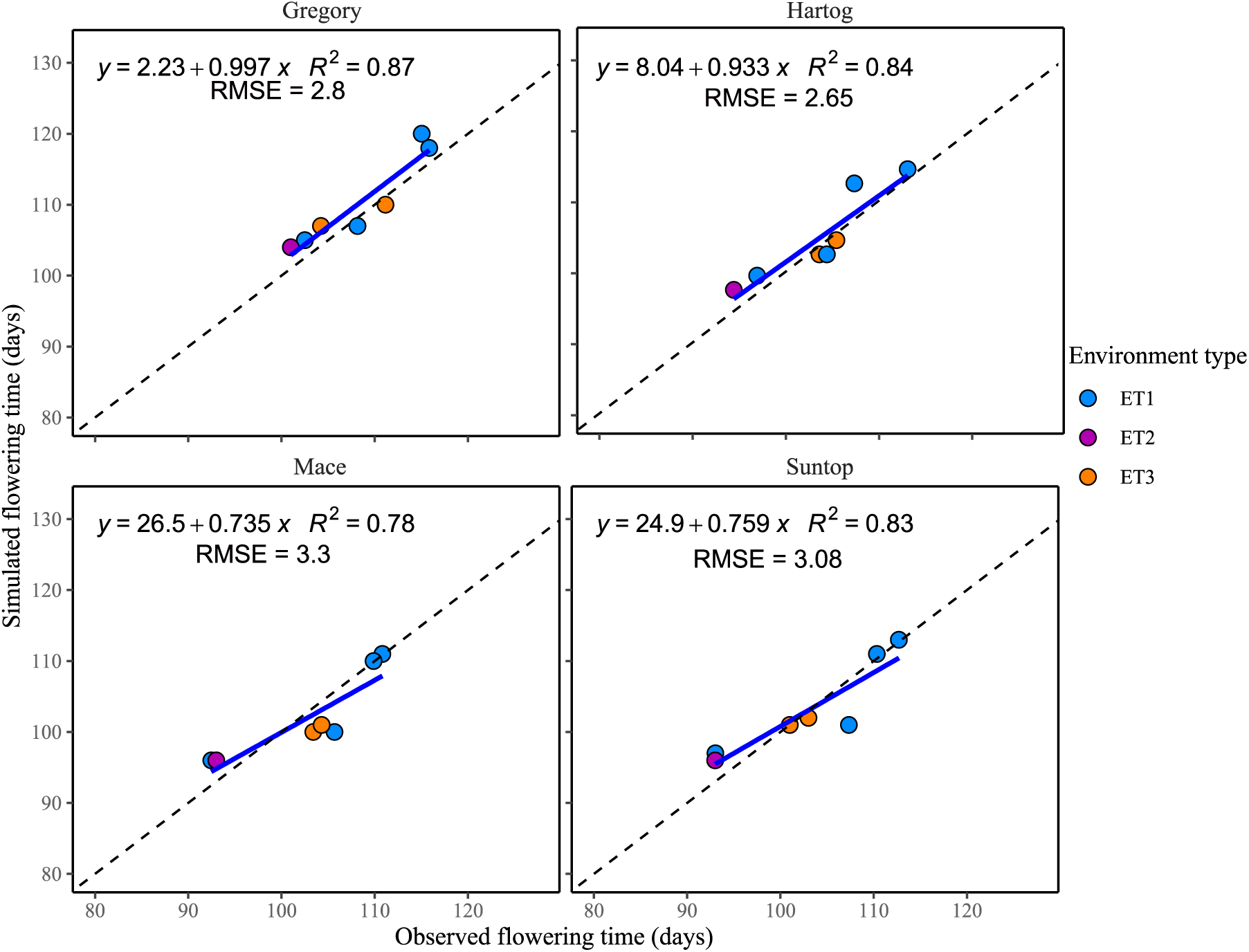
Flowering time simulated with the crop model against observed mean values for four parental genotypes in the studied trials. Flowering time is expressed in days after sowing. To best simulate the growth and development of the tested cultivars at each trial, an APSIM parameter (thermal time to floral initiation, ‘tt_floral_initiation’) was tuned so that simulated flowering time matched observed flowering time. For each genotype, the fitted relationship is presented as a solid blue line, and the 1:1 line is presented as a dashed black line.

**Fig. S2.**
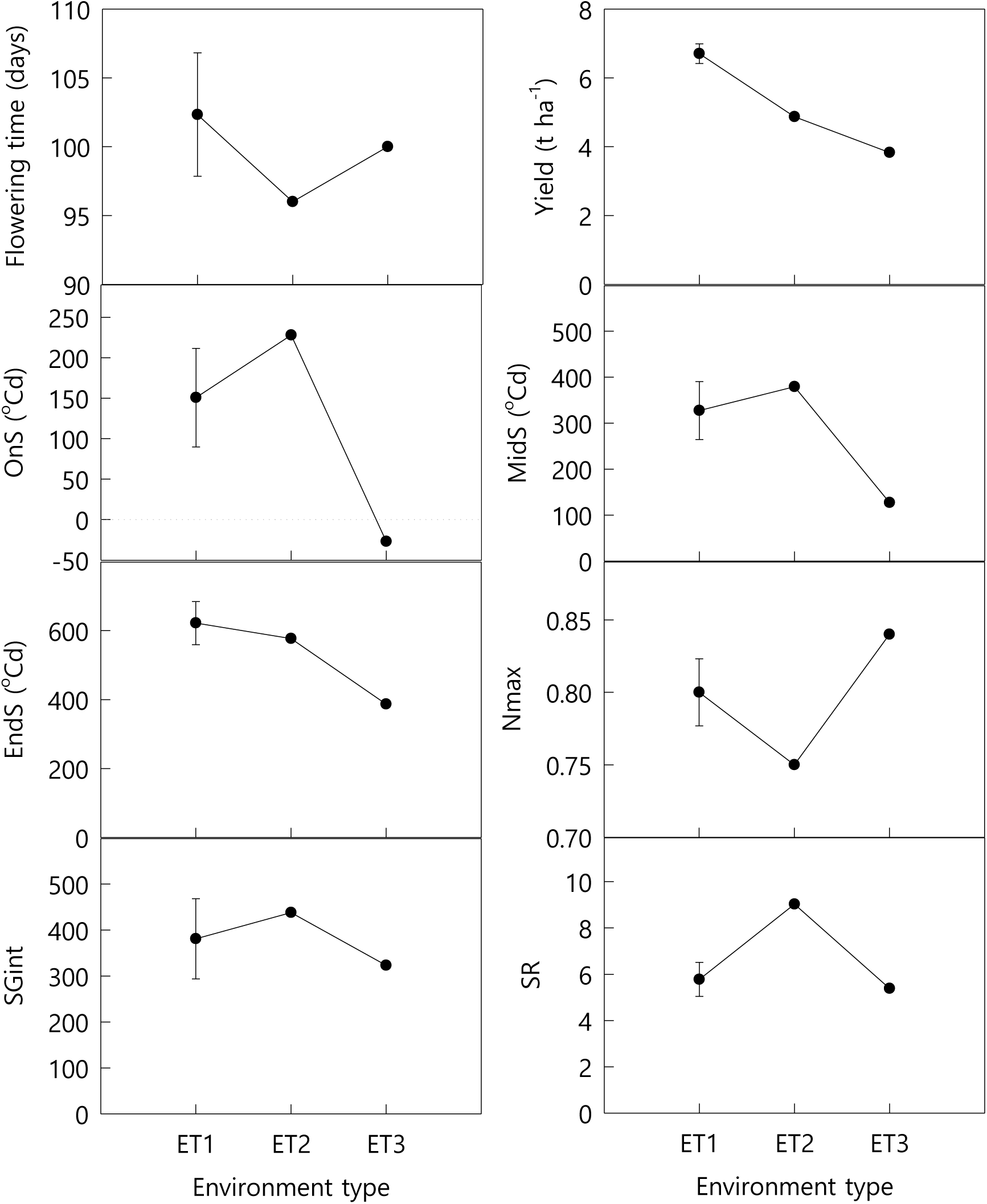
Average yield, flowering time, and stay-green traits in the different environment types for the five trials where stay-green traits were measured. Stay-green traits are start of senescence (OnS, °Cd since flowering), mid of senescence (MidS, °Cd since flowering), end of senescence (EndS, °Cd since flowering), indicator of the rate of senescence (SR), maximum greenness (Nmax), stay-green integral (SGint). Colors correspond to the environment types (ETs). Bars correspond to standard error.

